# OmniAge: a compendium of aging omic biomarkers links mitotic clocks to clonal hematopoiesis and causality

**DOI:** 10.64898/2026.04.29.720033

**Authors:** Zhaozhen Du, Yunchao Ling, Huige Tong, Xiaolong Guo, Andrew E. Teschendorff

## Abstract

Interest in aging ‘omic’ biomarkers has grown due to their ability to quantify biological age. Most of these biomarkers have been derived in blood and fall into many diverse categories, yet relatively little is known about their correlative patterns, especially between biomarkers from different categories. Here we present the *OmniAge* R and Python package, a collection of 413 aging omic biomarkers representing 12 different categories, including traditional epigenetic clocks, epigenetic mitotic clocks, DNA methylation-based proxies for clonal hematopoiesis and inflammaging, causal clocks, cell-type specific epigenetic clocks and single-cell transcriptomic clocks. By studying their inter-class correlations across large blood datasets, we reveal associations of mitotic age with clonal hematopoiesis subtypes and causal clocks, which are predictive of cancer risk. Using proxies of serum protein levels, we further dissect associations with mitotic clocks, clonal hematopoiesis and causal clocks into distinct biological processes mapping to key aging pathways. Applying *OmniAge* to multi-modal data of sorted immune cell-types reveals that age-acceleration derived from transcriptomic and epigenetic clocks correlate, but that this is driven by underlying cell-type heterogeneity. In summary, the *OmniAge* package is an exploratory tool for evaluating large numbers of aging omic biomarkers, and to aid discovery and generate new hypotheses.

---

Aging biomarkers hold great promise to help triage individuals in a population to treatments or lifestyle interventions that can prevent or delay disease onset, hence improving their health-and-lifespan ^1^. Aging ‘omic’ biomarkers (AOB) ^2–5^ have attracted strong interest, not only because they have been reproducibly associated with quantifiable measures of biological age ^6^ such as all-cause mortality ^7–9^, but also because the molecular alterations underpinning these markers may be indirectly or directly causative of accelerated aging ^10,11^. AOBs display a remarkably diversity: they comprise traditional DNA methylation (DNAm)-based clocks that include 1^st^ generation (trained on chronological age) ^12–14^, 2^nd^ generation (trained on clinical biomarkers, exposures, health outcomes) ^8,9,15^ and 3^rd^ generation (trained on longitudinal changes in clinical aging biomarkers) ^6,16^ ones, DNAm-clocks that yield proxies of cellular age, including telomere length ^17^ and mitotic age ^18–22^, cell-type specific DNAm-clocks ^23^ and transcriptomic clocks built from single-cell RNA-Seq data ^24–28^ that yield proxies of biological age at cell-type resolution ^23,29^, as well as omic surrogate predictors of serum protein markers and clonal hematopoiesis ^30–35^. In addition, AOBs have also been built from other omic data-types including proteomic data ^2,36–38^.

In view of this diversity, it becomes important to collate them all into one common tool, to facilitate comparative analyses. Since different AOBs may capture different aspects of aging biology, such comparative or cross-correlation analyses could shed important novel insights into the molecular aging process. For instance, non-trivial associations between AOBs could point to important connections between different aging processes or hallmarks. Whilst a number of distinct tools that facilitate such inter-AOB comparisons have been produced ^39–41^, these often omit or underrepresent important categories of AOBs, which can limit the discovery of novel insights. To address this shortcoming, we here present the *OmniAge* R and Python package, a simple user-friendly software tool, encompassing 413 AOBs across 12 diverse categories to help facilitate novel discoveries. As such, *OmniAge* comprises significantly more AOBs than competing tools.

As proof-of-principle, we apply *OmniAge* to a large series of whole blood DNAm datasets, revealing previously unknown associations between serum protein level proxies, mitotic clocks, clonal hematopoiesis and causal clocks. Using one of the largest prospective cancer studies, we further demonstrate that these associations lead to important novel insights for blood-based cancer risk prediction. With *OmniAge*, we also demonstrate that molecular clock age-acceleration estimates derived from different data modalities (transcriptomics & DNA methylation) can correlate, although driven by underlying cell-type heterogeneity. The findings we present here hinge on the much larger compendium of AOBs in *OmniAge* and would have been missed with other existing tools.

## Results

### *OmniAge*: a more diverse aging omic biomarker toolkit

Overall, in the *OmniAge* tool, we assembled a total of 413 AOBs representing 12 different categories (**Fig.1a**, **SI table S1**). Compared to other tools like *pyaging* ^39^, *methylCIPHER* ^40^ and *BioLearn* ^41^, *OmniAge* includes significantly more AOBs, as well as a more diverse set of AOBs (**Fig.1b**, **SI table S1**). For instance, *OmniAge* includes many DNAm-based mitotic clocks in the cellular aging clock category, which are absent from most other tools (**Fig.1b**). These mitotic clocks yield proxies for the mitotic age (i.e. the age-cumulative number that stem-cell and amplifying progenitors have undergone) and have been shown to be particularly powerful for predicting cancer risk in solid tissues ^18,20,22^, although their assessment in blood tissue remains largely unexplored. Our software also includes recently published DNAm-based proxies (surrogate markers) for hsCRP-levels (inflammaging) ^30,31^, various types of clonal hematopoiesis (CH) ^35^ and serum protein levels (Episcores) ^33^ (**Fig.1b**). Unlike other tools, we also include some of the recently published cell-type specific clocks, including both DNAm-based ones ^23^ and scRNA-sequencing based ^24^ (**Fig.1b**). As we show further below, the substantial increase of AOBs in certain AOB-classes can help identify previously unknown associations between different types of AOBs. To ensure the quality of the resource, we tested each AOB within the resource, comparing the output of each one to the one obtained from the original software derived from the original publication, finding in each instance a perfect correlation (**SI figs.S1-S2**).

**Figure-1.**
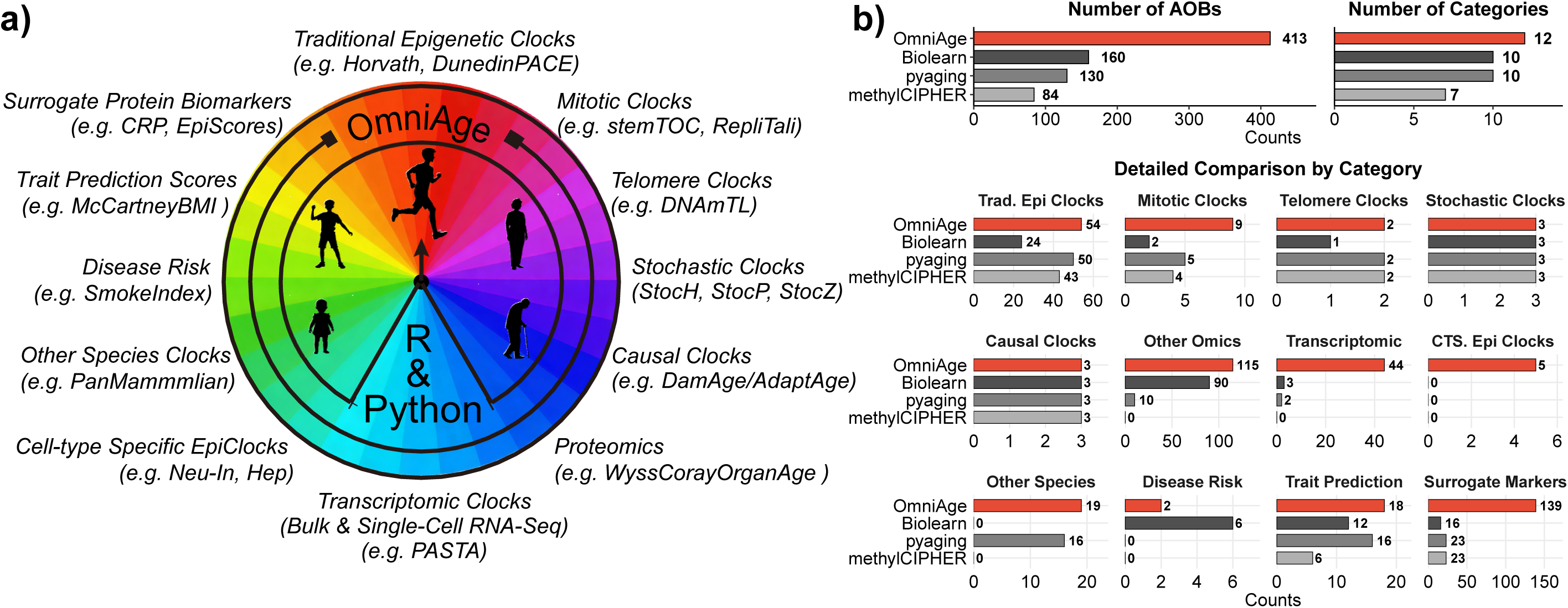
Graphical summary of the OmniAge R/Python package. **a)** The logo illustrates the diversity of OmniAge, encompassing a wide variety of aging omic biomarkers (AOBs), and integrating them in both R and Python environments. **b)** Benchmarking OmniAge against other aging biomarker tools. Top left panel: Comparison of the total number of supported AOBs across OmniAge, pyaging, Biolearn and methylCIPHER. Top right panel: Barplot compares the number of diverse AOB categories per tool. Bottom panel: Distribution of AOBs across various categories. Counts for each tool are represented by colored bars as indicated in the legend.

### Supervised and unsupervised clustering of AOBs reveal robust patterns of association

We first explored the overall correlation landscape between human DNAm-based AOBs in whole blood using one of the largest available cohorts encompassing 729 White Caucasian samples (Johansson cohort, Illumina 450k, **Methods, SI table S2, SI table S3**) ^42^. Initially, we stratified the AOBs according to their defined categories and, for visualization purposes, restricted analysis to a subset of around 40 AOBs. Estimating sex and age adjusted age-accelerations for each AOB, we found that AOBs within a category displayed stronger correlations compared to AOBs across categories (**SI fig.S3**). These relative correlation patterns were replicated in the Illumina 450k Hannum cohort comprising Hispanics in addition to White Caucasians ^43^, and the TZH cohort, generated with EPICv1 and comprising Han Chinese ^44^ (**SI table S2**, **SI fig.S3, SI tables S4-S5**). To explore the inter-AOB correlations in more detail, we next performed PCA-analysis over all human DNAm-based AOBs. This also revealed broad consistency between the 3 cohorts (**SI fig.S4**). For instance, whilst surrogate markers displayed a big spread, in each of the 3 cohorts, PC1 distinguished mitotic and CH-proxies from traditional epigenetic clocks, whilst PC2 distinguished trait-predictors (e.g BMI), intrinsic inflammatory markers (e.g intCRP) and biological aging clocks (e.g. DunedinPACE) from chronological aging clocks (**SI fig.S4**). Performing unsupervised clustering in the Johansson cohort using the focused subset of 40 AOBs (**Methods**), revealed a total of 8 AOB clusters, which broadly speaking were correlated with the defined AOB categories (**Fig.2a**). For instance, AOBs of CH and mitotic age formed respective tight groups, but interestingly, the DNMT3A clonal hematopoiesis AOB correlated more strongly with mitotic clocks than with TET2 or ASXL1 AOBs. Another tight cluster was formed by AOBs representing biological aging clocks, including notably DunedinPACE (aimed at estimating the pace of aging) ^6^, GrimAge (clock that associates strongly with smoking exposure and mortality) ^8,9^ and Systems Age (a clock that encapsulates aging across 11 different tissues) ^45^ clocks. The only AOB missing from this cluster was PhenoAge ^15^ and its PC-version ^46^, which both mapped to the biggest cluster composed of diverse AOB categories including chronological aging clocks and causal clocks. Of note, AdaptAge (a clock constructed from putative causal CpGs associated with good health outcomes) did not cluster with DamAge (also constructed from putative causal CpGs but associated with adverse health outcomes) ^11^, owing to their strong anticorrelation. Thus, in terms of absolute correlation, all causality clocks formed one tight group. Interestingly, another cluster mapped more closely to chronological aging clocks, which included the most accurate clock for chronological aging (Zhang clock ^12^) as well as its stochastic version ^47^. Although clocks for telomere length displayed strongest correlations with the other cellular aging clock category, i.e mitotic clocks, they still formed a separate group. Importantly, most of these AOB clusters were replicated in the Hannum and TZH whole blood cohorts **(Fig.2a-b, Methods**), despite differences in ethnicity and platform. For instance, the CH, mitotic, poor health outcome, telomere and inflammaging (CRP) clusters were all well replicated. The chronological aging AOB cluster was well replicated in one of the cohorts (Hannum) but displayed overlap with the main AOB-cluster in TZH (**Fig.2b**). To further confirm this, we compared the unsupervised clustering result obtained in another large whole blood cohort of White Caucasians (the Airwave study) performed with EPICv1 arrays to Johansson (Whites+450k) and TZH (Han Chinese+EPICv1), revealing strong concordance and high Adjusted Rand Indices (**Methods, SI fig.S5**). Overall, these data show that the relationship between the various AOBs is robust across ethnicities (White Caucasian; Han Chinese; Hispanics) and platforms (EPICv1 vs 450k).

**Figure-2.**
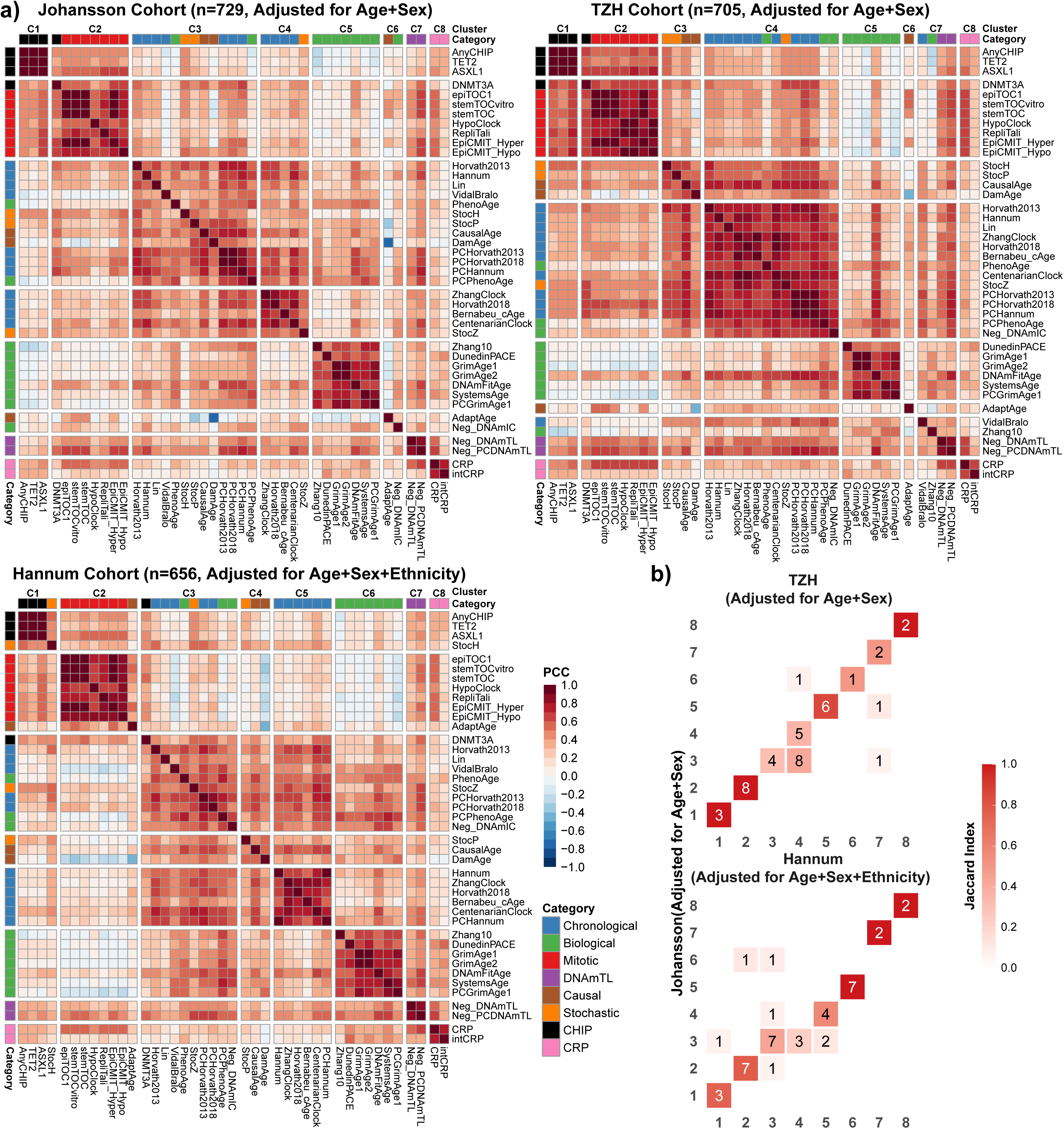
Unsupervised clustering of Aging Omics Biomarkers (AOBs). **a)** Heatmaps display pairwise Pearson correlations of selected AOBs (their residuals after adjusting for age, sex and if necessary also ethnicity) in the Johansson (n=729), TZH (n=705), and Hannum (n=656) cohorts. AOB residuals were clustered using a Gaussian mixture EM-algorithm. For the Johansson cohort, the optimal number of clusters (k = 8) was selected using the BIC, whereas the clustering for the TZH, and Hannum cohorts was explicitly constrained to an 8-cluster solution (k = 8) to ensure structural comparability. The top annotation bar indicates the assigned cluster (e.g. C1, C2), while the left annotation bar represents the defined functional category of each biomarker. The color scale indicates the Pearson correlation coefficient (red = positive, blue = negative). **b)** Confusion matrices illustrating the concordance of cluster assignments between the Johansson (y-axis) and TZH (x-axis) and Hannum (x-axis) cohorts. Tile colors indicate the Jaccard Index (defined as the ratio of shared biomarkers to the total number of unique biomarkers in the pair), while the number within each tile represents the absolute count of overlapping biomarkers.

### Mitotic and stochastic clocks associate with clonal hematopoiesis

The collation and analysis of a diverse set of AOBs can lead to novel biological insights. For instance, we asked which AOBs demonstrate strongest association with clonal hematopoiesis (CH) proxies, as currently it is unknown which AOBs (if any) correlate with CH. We observed that mitotic clocks in particular displayed the strongest and most consistent associations with CH-proxies ^35^ (**Fig.3a-b**). This is consistent with mitotic clocks ‘tracking’ the proliferative clonal expansions that define CH.

**Figure-3.**
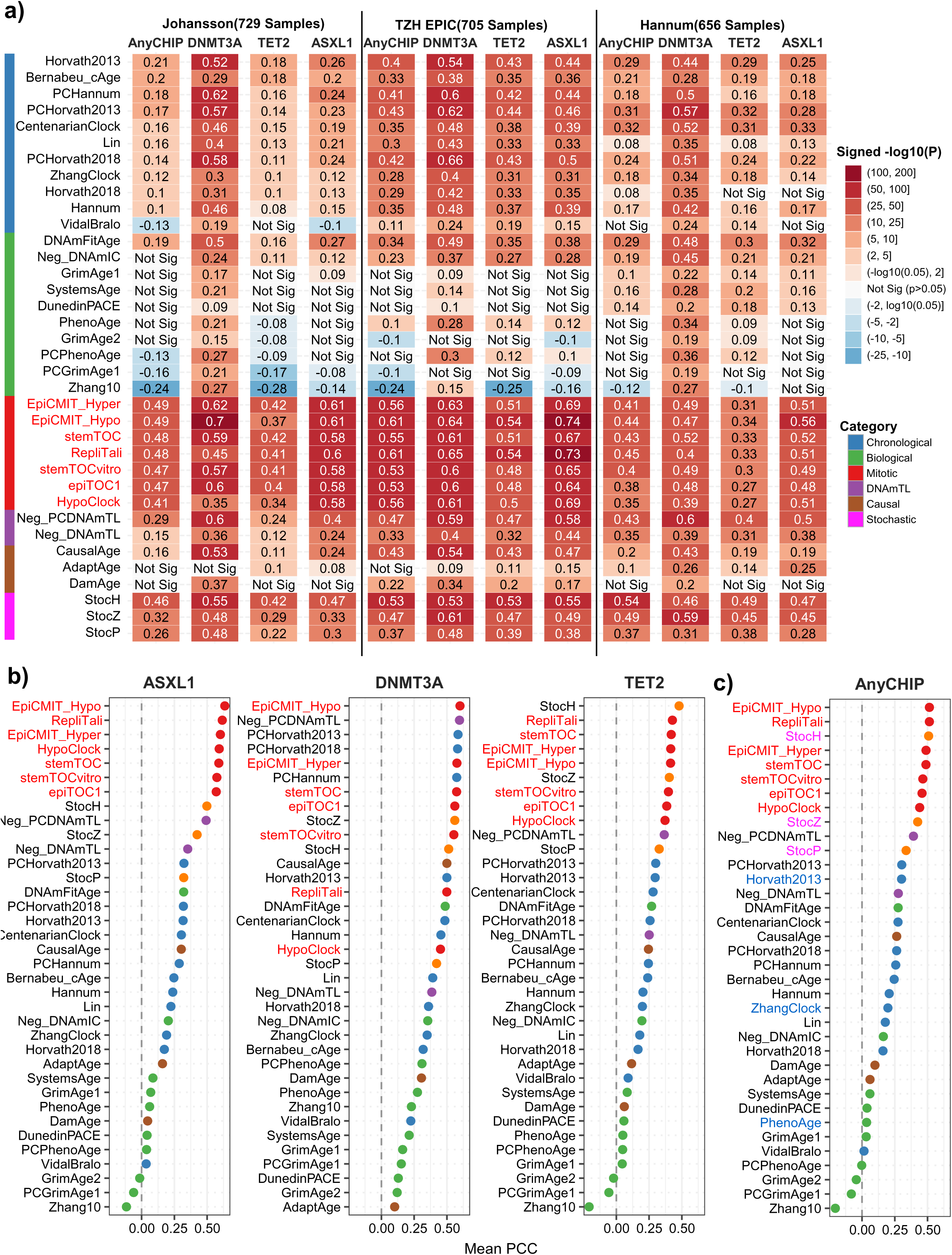
Associations of epigenetic clocks with Clonal Hematopoietic Proxies. **a)** Heatmap displaying the Pearson correlation coefficients (PCC) between epigenetic age acceleration and CHIP status in the Johansson (n=729), TZH EPIC (n=705), and Hannum (n=656) cohorts. Correlations were calculated using residuals adjusted for age and sex (and ethnicity for the Hannum cohort). Rows represent specific epigenetic clocks, annotated with a vertical color bar that indicates their functional category. Columns represent CHIP subtypes, including “AnyCHIP” (presence of a mutation in any of one of DNMT3A, TET2 or ASXL1. Tile colors indicate the signed -log10(P-value), reflecting both the direction (red for positive, blue for negative) and statistical significance of the association. Numerical values within tiles denote the PCC; non-significant associations (P > 0.05) are labeled “Not Sig”. P-values less than 1e-5 are significant under a strict Bonferroni-adjustment. **b)** Dotplots displaying the mean PCC for each epigenetic clock, as averaged across all three cohorts, and with clocks ranked by decreasing mean correlation value. Points are colored according to the functional category of the epigenetic clock. **c)** As b), but for the AnyCHIP proxy.

Other interesting associations with CH were displayed by the stochastic clocks ^47^. These stochastic clocks displayed stronger associations with CH than the clocks they derive from (**Fig.3c**). For instance, the stochastic version of Horvath’s clock (StocH) correlated more strongly with the CH-proxy than the Horvath clock itself (**Fig.3c**). Whilst the CpGs making up the stochastic versions of Horvath, PhenoAge and Zhang clocks are identical to those making up the corresponding Horvath, PhenoAge and Zhang clocks, the stronger associations with CH suggest that the underlying DNAm changes caused by *TET2/DNMT3A/ASXL1* mutations are of an intrinsically more stochastic nature.

### Mitotic clocks associate with AdaptAge causal clock

Besides CH, mitotic clocks also displayed a surprisingly strong positive association with AdaptAge, yet interestingly not with DamAge (**Fig.4a**). The association with AdaptAge was however only seen for mitotic clocks based on age-associated hypermethylation (e.g stemTOC, epiTOC1/2), and were not consistent for those based on hypomethylation (e.g. epiCMIT-hypo or RepliTali) (**Fig.4a**). Interestingly, the association between stemTOC and AdaptAge was age-dependent, with the correlation being much stronger in the older (i.e. >60 years) age group compared to the young (<40 years) or middle-aged (40-60 years) ones (**Fig.4b-c, SI fig.S6**). Thus, the correlation of stemTOC with CH and AdaptAge, specially in the older age-groups, suggests that mitotic clocks like stemTOC may be capturing beneficial clonal expansions. In relation to this, it is worth noting that although CH has generally been associated with adverse health outcomes ^48–50^, some studies have indicated beneficial clonal mosaicism, for instance in relation to Alzheimer’s Disease ^51^. Of note, both AdaptAge and DamAge correlated poorly with DNMT3A, TET2 and ASXL1-associated CH, consistently ranking in the bottom half of AOBs (**Fig.3b-c**). Thus, despite stemTOC correlating strongly with both CH and AdaptAge, CH associated with mutations in epigenetic enzymes did not correlate with any of the causal clocks. This may indicate that characterizing CH merely by mutations in DNMT3A, TET2 or ASXL1 may capture both beneficial and damaging clonal expansions.

**Figure-4.**
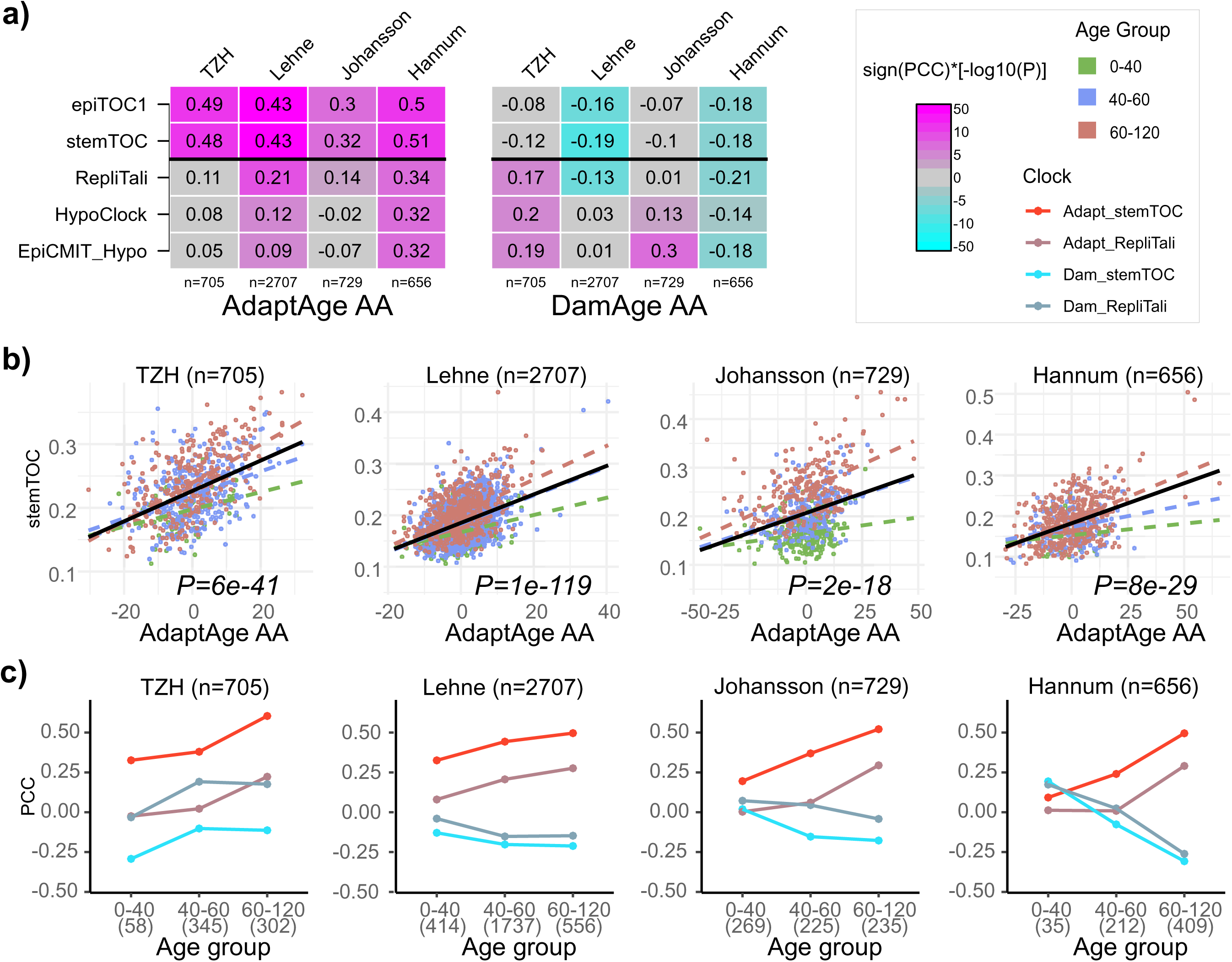
Age-dependent correlation of mitotic clocks with AdaptAge. **a)** Heatmaps of Pearson Correlation Coefficients (PCC) between 5 mitotic clocks with AdaptAge and DamAge age-acceleration (AA) across 4 large whole blood cohorts. Sample size of cohort is given. Colors in heatmap have been labeled according to signed statistical significance, as indicated in figure legends box. The top two mitotic clocks are based on gains of DNAm with age, the bottom 3 are based on loss of DNAm with age. The entries in the heatmaps are the PCCs. **b)** Scatterplot of stemTOC’s mitotic age estimate (y-axis) against the age-acceleration (AA, x-axis) from the AdaptAge clock for 4 different DNAm datasets, with samples colored by age-group. Regression fit (black) and P-value from a linear regression fit to all samples is given. Also shown are the regression lines within each age-group. **c)** Plots displaying the Pearson Correlation Coefficient of stemTOC with AdaptAgeAA and DamAgeAA, and of RepliTali with AdaptAgeAA and DamAgeAA, as a function of age-group.

### *OmniAge* identifies complex patterns of association with cancer risk

The strong correlation between mitotic clocks like stemTOC or epiTOC1 with AdaptAge was a surprising finding. To study its biological significance, we thus applied *OmniAge* to Illumina 450k DNAm data of the European Prospective Investigation Cancer Italy cohort (EPIC-Italy), which is a prospective study in which 845 whole blood samples (after QC, **Methods, SI table 6**) were taken years before eventual diagnosis of cancer ^52,53^. Using all available samples we first verified the previously identified patterns: mitotic clocks like stemTOC and epiTOC1 correlated strongly with AdaptAge but negatively with DamAge, whilst also displaying strong positive correlations with CH-proxies (**Fig.5a**). Consistent with the earlier analysis, CH-proxies generally did not correlate well with either AdaptAge or DamAge, except for ASXL1-CH (**Fig.5b**). We next performed univariate and multivariate Cox-regression analyses on two case-control sub-cohorts, where individuals developed either breast cancer (females only, n=233 cases and n=340 controls) or colon cancer (both sexes, n=139 cases and n=424 controls) (**Methods**). In the case of breast cancer, mitotic clocks and CH-proxies displayed the most pronounced increased hazard ratios, which remained significant after adjustment for age, immune cell fractions and multiple testing (**SI fig.S7-S8, SI table S7**). Importantly, AdaptAge, stemTOC and TET2/ASXL1 CH-proxies were all significantly associated with poor outcome, even after adjustment for age and multiple testing (**Fig.5c, SI table S7**). When additionally adjusting for immune-cell fractions, hazard ratios remained largely invariant, although P-values became more marginal due to loss of power (**SI table S7**). In the case of colon cancer, the pattern was very different: here DamAge and DNMT3A-CH increased risk of cancer-diagnosis whilst AdaptAge decreased cancer-risk, associations which remained significant upon adjustment for multiple testing, age and sex (**SI table S8, Fig.5d**). In the case of colon-cancer, neither stemTOC nor TET2/ASXL1 displayed significant associations (**Fig.5d, SI table S8**). Because the naïve CD4T-cell fraction did display a strong protective effect, when adjusting for IC-fractions, only the associations for DamAge and AdaptAge remained significant. Overall, these data suggest that the CH-proxies in the EPIC-Italy cohort are generally associated with increased risk of cancer, albeit with different CH-subtypes associating with different cancer-types, and importantly that AdaptAge does not always associate with good health outcomes. Thus, as illustrated in the case of breast cancer, the positive correlation of a mitotic clock like stemTOC with breast cancer diagnosis and TET2/ASXL1 CH, points towards the mitotic clock capturing CH with an adverse outcome.

**Figure-5:**
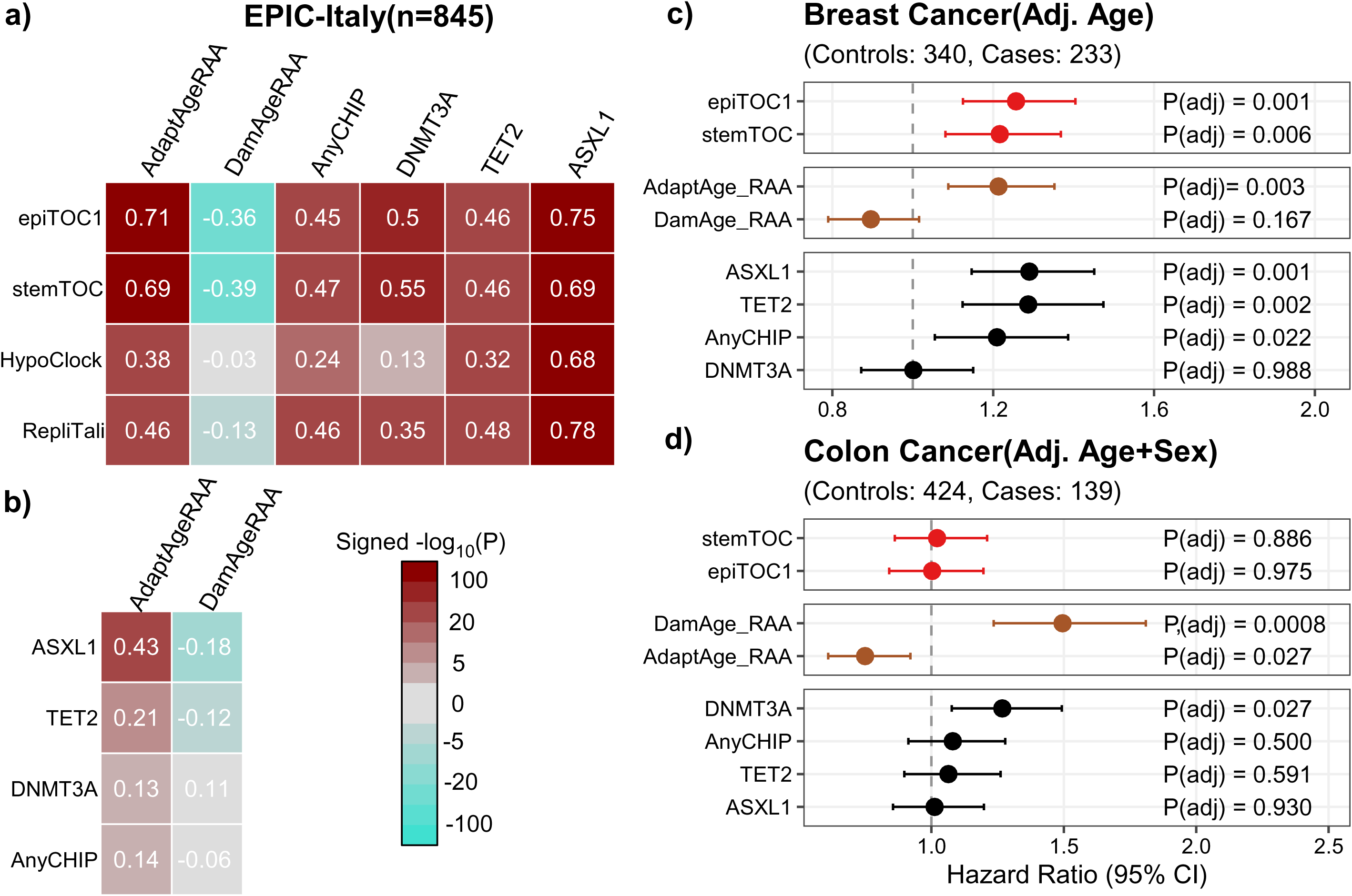
Mitotic clocks and clonal hematopoiesis associations with cancer risk. **a)** Correlation heatmap of 4 selected mitotic clocks with the residual age-acceleration (RAA) of AdaptAge, DamAge and 4 clonal hematopoiesis (CH) proxies, as calculated using all available blood samples from EPIC-Italy cohort. Values in heatmap indicate the Pearson Correlation Coefficient (PCC). Colors indicate signed statistical significance: sign(PCC)*[-log_10_P]. **b)** As a), but displaying the PCCs of CH-proxies with AdaptAge-RAA and DamAge-RAA. **c)** Hazard Ratio plots in EPIC-Italy restricting to a prospective breast cancer case – control subcohort (females only). Number of cases and controls is indicated. Plots indicate the Hazard Ratio (HR), lower and upper 95% confidence intervals, as derived from a Cox-regression, adjusting for age. P-values are from the Wald test and were adjusted using the Benjamini-Hochberg (BH) method. **d)** As c) but for the prospective colon cancer subcohort, where adjustment in the Cox-regression also included a covariate for sex.

### *OmniAge* reveals associations of serum protein proxies with mitotic clocks and CH

To better understand the mechanistic underpinnings of the previous associations, we applied *OmniAge* to correlate DNAm-proxies of 109 serum protein levels (Episcores)^33^ to mitotic-clocks, casual clocks and our proxies for clonal hematopoiesis. We supplemented the Episcores with more accurate proxies for high sensitive serum C-reactive protein levels (CRP)^54^ and interleukin-6 (IL6) ^32^. Unsupervised clustering of Pearson correlations between Episcores and the other AOBs, as assessed in the same EPIC-Italy cohort, revealed the presence of 3 broad clusters (**Fig.6a**). One cluster (magenta cluster, **Fig.6a**) was marked by high positive correlations of mitotic clocks and the ASXL1-CH proxy with Episcores, which included markers of chronic inflammation (e.g. CRP, TNFRSF1B), of increased cytotoxic lymphocyte (NK/T-cell) activity (e.g. GZMA, Granulysin, CD48, and LY9), of chemokines involved in T-cell trafficking (CCL11, CCL17, CCL18, CCL21, CCL22 and CXCL10/11), of extracellular matrix degradation and fibrosis (MMP-1, MMP-2, MMP-12, FAP, THBS2), of vascular remodeling (ESM-1, Semaphorin-3E), of metabolic and hormone dysregulation (GHR, Insulin-receptor, FGF21 and RARRES2) and finally also of hematopoietic stem-cell niche dysregulation (TPO, G-CSF and EZR). In contrast, a cluster displaying an opposite trend (negative correlations) (blue-cluster, **Fig.6a**) included markers of neutrophil-dominated acute inflammation (Myeloperoxidase-MPO, Calprotectin-S100A9, and EN-RAGE-S100A12), as well as tissue-specific factors like hepatocyte-growth-factor activator (HGFA, liver), osteomodulin (bone) and neural adhesion molecule Contactin-4 (brain). We verified that the signed statistical significance levels in EPIC-Italy were congruent with those obtained in another blood cohort (Johansson) (**SI fig.S9**), and that unsupervised clustering in the Johansson cohort led to a similar grouping as in EPIC-Italy (**SI fig.S10**). Ranking all Episcores according to their absolute correlation strength with the other aforementioned AOBs, adjusted for age and sex, confirmed these correlative patterns with mitotic clocks and CH-proxies (**Fig.6b**).

**Figure-6:**
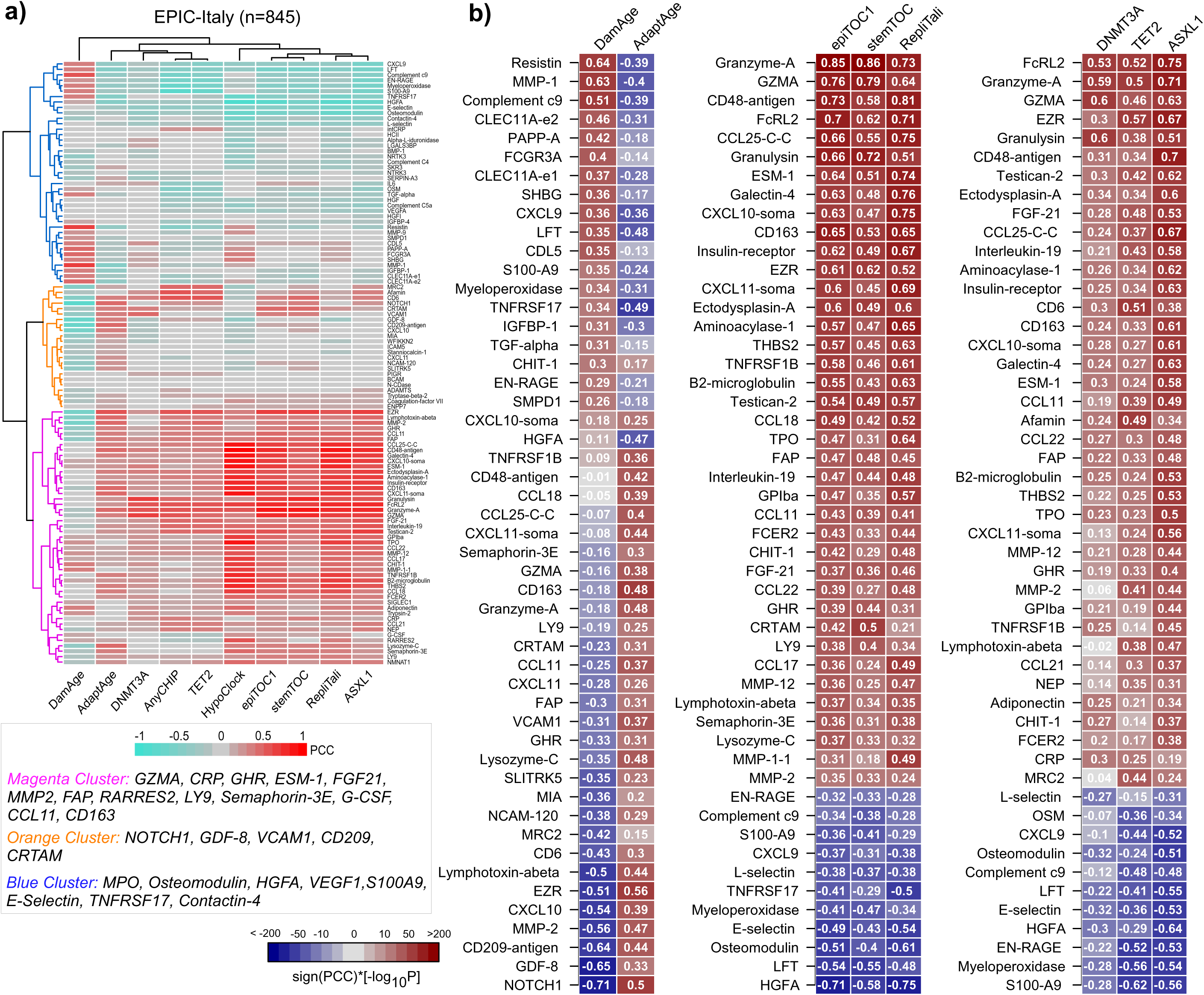
Correlations of serum protein proxies with CH, mitotic and causal clocks. **a)** Unsupervised clustering heatmap using Euclidean distance metric of the Pearson correlation coefficients (PCC) between 111 Episcores (DNAm-based proxies of 111 serum protein levels) and 10 other AOBs including DamAge, AdaptAge, 4 mitotic clocks (stemTOC, epiTOC1, RepliTali, HypoClock), and the 4 CH-proxies. Three main clusters are labeled with distinct colors and examples of proteins in each cluster are shown below heatmap. PCCs between Episcores and the other 10 AOBs were adjusted for age and sex. **b)** Heatmaps displaying the signed statistical significance levels of the PCCs for the top-50 Episcores correlating with causal clocks, 3 mitotic clocks (stemTOC, epiTOC1 and RepliTali) (HypoClock was excluded because it is confounded with variations in immune cell fractions) and the 4 CH-proxies. Episcores have been ranked according to directionality of the correlations. The numerical entries in the heatmap are the PCCs. P-values less than 1e-5 are significant under a strict Bonferroni-adjustment.

Age and sex adjusted correlations of Episcores with the causal clocks also revealed a clear striking dichotomy: among the most strongly correlated Episcores, those correlating positively with DamAge, displayed the exact opposite trend with AdaptAge and vice-versa (**Fig.6b**), in line with the opposing roles of these two clocks. For instance, the three Episcores correlating most negatively with DamAge were NOTCH1, GDF-8 and antigen CD209, all of which correlated positively with AdaptAge (**Fig.6b**). Interestingly, NOTCH1 is known to promote tissue-repair, regeneration and to maintain stem and progenitor cell pools, with a decline in NOTCH1 signaling having been associated with age-related diseases ^55–57^. GDF-8 (Myostatin) is a TGF-β superfamily member that negatively regulates skeletal muscle mass.

The observed anti-correlation with a DNAm-clock like DamAge could reflect complex dysregulation of the muscle-niche environment ^58–60^. CD209 (a C-type lectin receptor) is expressed on dendritic cells and monocytes and is a known sentinel of immune-surveillance. The observed anti-correlation with DamAge would be consistent with a dysfunctional dendritic cell compartment being a feature of a high-damage state.

Overall, these associations of Episcores with mitotic clocks, CH-proxies and causal clocks highlight diverse biological processes (e.g. inflammation, metabolic and hormone dysregulation, dysregulations of stem-cell niche, fibrosis and ECM degradation) that are associated with clonal expansions, whilst also dissecting them into distinct components, such as those associated with chronic low-grade inflammation and acute inflammation.

### An immune cell-type fraction clock

In the *OmniAge* package we also present a new class of molecular aging clock which tracks age-associated changes in immune cell type composition. The rationale for such a clock is as follows: because individual immune cell fractions (e.g. naïve CD4+ T-cells) have been shown to correlate with age and poor health outcomes ^61^, a composite measure that considers the whole immune cell-type composition of a sample could also be a potential aging biomarker (**SI fig.S11a**). To construct the clock, we used the NSPT cohort (EPIC arrays) of 3501 whole samples ^62^, estimating fractions for 12 immune cell-types using EpiDISH ^61,63^, subsequently performing model training and parameter optimization through an internal ten-fold cross-validation (**SI fig.S11a**). We considered two different machine learning algorithms (Random Forests + Lasso) to construct corresponding clocks for chronological age (**Methods**). Validating the clocks in 4 independent cohorts (Johansson et al (450k, n=729) ^42^, Hannum et al (450k, n=656) ^43^, Airwave (EPIC, n=1032**, SI table S2**) ^64^ and HPT (n=1394, **SI table S2**), revealed that the RandomForest (RF) algorithm obtained a higher validation accuracy (**SI fig.S12, SI fig.S11a-b**). Hence, we restricted subsequent analyses to this RF clock. Correlation of the age-acceleration from the RF immune-cell fraction clock with all AOBs (**SI table S9**) revealed particularly strong associations with the DNAm-based proxies of high sensitivity C-reactive protein (hsCRP) levels, a marker of inflammaging (**SI fig.S11c**). Importantly, these associations largely disappeared when we restricted to the intrinsic hsCRP-score (**SI fig.S13a, Methods**), which by definition is adjusted for variations in IC-fractions. To demonstrate that our RF IC-fraction clock can be used to track disease-associated age-acceleration, we applied it to an EWAS DNAm dataset of Rheumatoid Arthritis (RA) ^65^, which confirmed age-acceleration in RA cases, driven mainly by a reduction in naïve CD8+ T-cells (**SI fig.S13b**).

### Cross-correlation of epigenetic and transcriptomic clocks

To further illustrate the functionality of *OmniAge*, we used it to address whether clock estimates from different data modalities are correlated or not. We assessed this in the MESA study ^66^, which profiled both DNAm (Illumina 450k) and mRNA expression (Illumina HumanHT-12 V4.0) in 1202 sorted classical monocytes and 214 sorted CD4+ T-cells (**Methods**). To estimate biological age in the transcriptomic data we applied single-cell CD4+ T-cell and monocyte clocks ^27^, as well as the multi-tissue multi-platform PASTA-clock ^26^, whilst for DNAm we considered Horvath’s pan-tissue clock, Zhang’s clock, Hannum’s clock and PhenoAge ^12,13,15,43^.

All DNAm and transcriptomic clocks validated well in the corresponding MESA CD4+ T-cell and monocyte DNAm and transcriptomic datasets, respectively (**SI fig.S14**). However, when we assessed the correlation in terms of the residual age-accelerations (adjusted for gender, race and cohort/lab), associations between the DNAm and single-cell transcriptomic RAAs were no longer significant, except for the transcriptomic CD4+ T-cell clock with Hannum in the MESA CD4+ T-cell samples, and the transcriptomic monocyte clock with PhenoAge in the MESA monocyte samples (**SI fig.S15a-b**). On the other hand, the RAA of the multi-platform transcriptomic PASTA clock displayed much stronger correlations with PhenoAge and Hannum clocks, although not for Horvath or Zhang, the latter two clocks being more aimed at predicting chronological age (**Fig.7a**). The highly significant correlations with Hannum and PhenoAge clocks, suggested to us that this is capturing a genuine signature of biological aging, of which an increase in the memory to naïve CD4+ T-cell subfractions is a natural candidate ^61^. To explore this, we applied PASTA to two large scRNA-Seq datasets ^67,68^, in order to compare PASTA-scores between naïve and memory CD4+ T-cell subsets from the same donors (thus age-matched) (**Methods**). Validating the applicability of PASTA to these scRNA-Seq datasets, PASTA scores were well-correlated with age in both memory and naïve subsets (Fig.7b), and confirming our hypothesis, PASTA-scores were indeed higher in the memory subset compared to naïve CD4+ T-cells (**Fig.7b**), a result that we were able to further validate on two sorted mRNA microarray datasets (**Fig.7c**). That DNAm-clocks predict higher DNAm-Ages in memory T-cells compared to naïve counterparts of same chronological age is well-known ^69^, and we were able to further confirm this by application of the DNAm clocks to the sorted EPIC DNAm dataset of Salas et al ^70^ (**Fig.7d**). Thus, the observed correlation between PASTA and Hannum/PhenoAge in the sorted CD4+ T-cells from the MESA study is likely driven by the well-known age-related shift of naïve CD4+ T-cell to memory CD4+ T-cell subfractions ^61^. Supporting this, we applied EpiDISH to the CD4T-cells to estimate the relative naïve and memory subfractions, subsequently adjusting the DNAmAge clock estimates for this variation, which led to a marked reduction in the correlations, rendering them not-significant (**Fig.7e**). In contrast to the CD4+ T-cells, correlations in monocytes were much weaker and not significant except for the PhenoAge clock (**SI fig.S15c)**. This suggests that the PhenoAge clock is capturing a process of biological aging in monocytes that is also reflected at the transcriptomic level, consistent with findings reported elsewhere ^30^. Overall, these results demonstrate that epigenetic and transcriptomic clocks both capture known age-related shifts in cell-type composition, and that once adjusted for this underlying cell-type heterogeneity, associations are much weaker.

**Figure-7:**
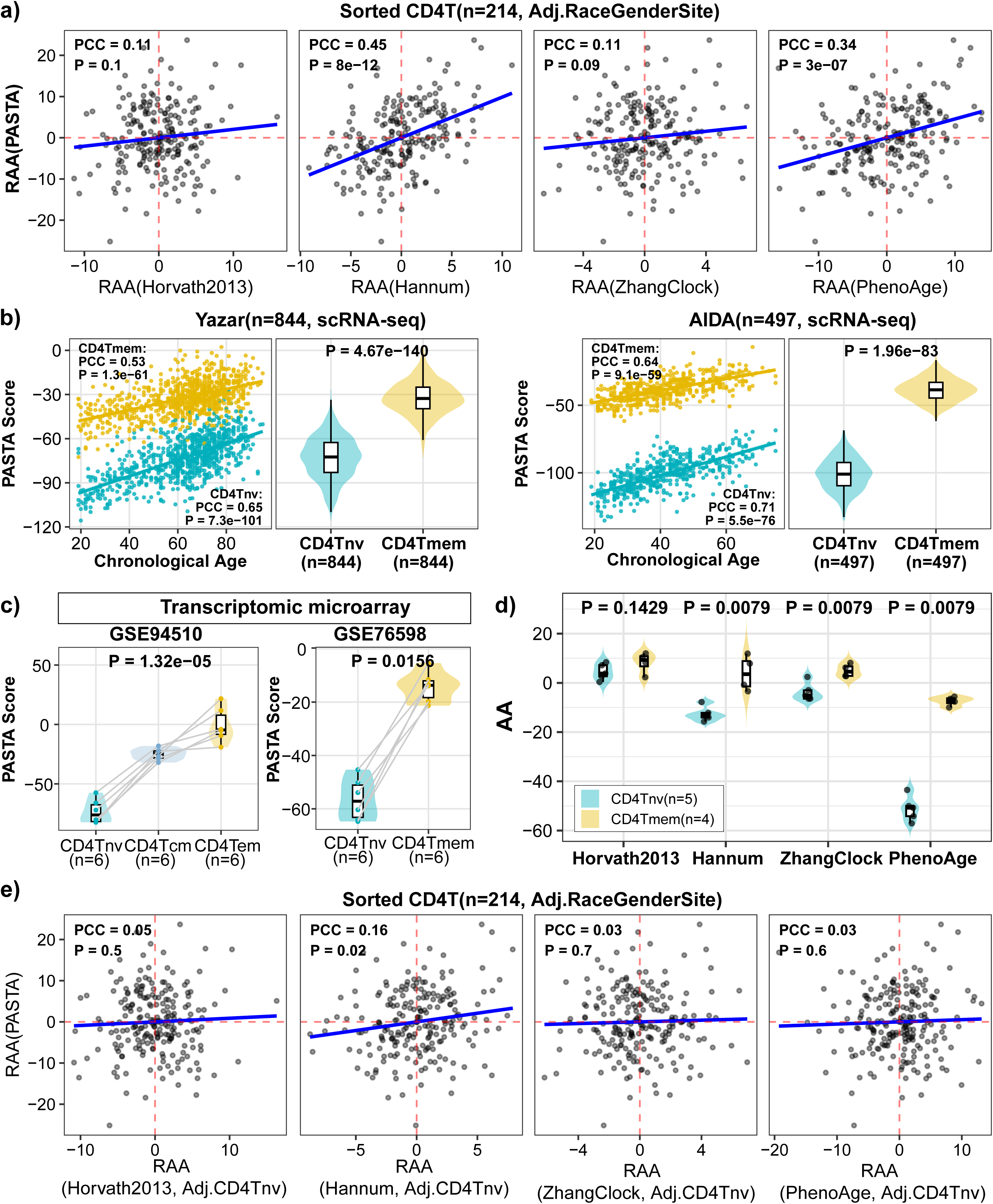
Cross-correlation of transcriptomic and epigenetic clocks in CD4+ T cells. **a)** Scatterplots display the relationship between Residual Age Acceleration (RAA) derived from the PASTA transcriptomic estimator and four established epigenetic clocks (Horvath2013, Hannum, ZhangClock, and PhenoAge) using the MESA sorted CD4+ T cell dataset (n = 214). RAA values were adjusted for a composite covariate of race, gender, and study site. Pearson Correlation Coefficients (PCC) and associated P-values are shown. **b)** PASTA scores calculated from pseudobulk profiles of naive (CD4Tnv) and memory (CD4Tmem) CD4+ T cells, restricted to donors with ≥100 cells in both subsets (Yazar (OneK1K), n = 844 donors; AIDA, n = 497 donors). Left: Scatter plots depict PASTA scores against Chronological Age. Individual data points, each representing a single donor, and corresponding subset-specific regression lines are colored yellow for CD4Tmem and turquoise for CD4Tnv. Pearson correlation coefficients (PCC) and P-values are indicated for each subset. Right: Violin plots illustrate the distribution of PASTA scores for each subset. Pairwise comparisons between subsets are annotated with P-values from a one-tailed paired Wilcoxon signed-rank test. **c)** As b), but for microarray expression data of sorted CD4+ T-cell subsets. GSE94510: Differences between CD4Tnv, CD4Tcm, and CD4Tem were assessed via a linear regression to account for inter-individual variability across the 6 donors. GSE76598: Statistical significance was assessed using a one-tailed paired Wilcoxon signed-rank test. Gray lines indicate paired samples from the same donor. **d)** Epigenetic Absolute Age Acceleration (AA) across T cell differentiation states. Violin plots show the distribution of AA derived from four epigenetic clocks in CD4Tnv(n = 5) and CD4Tmem(n = 4) samples from the Salas sorted immune-cell DNAm dataset. P-values were calculated using a one-tailed Wilcoxon rank-sum test. **e)** As a), but when adjusting the DNAm-clock DNAmAge estimates also for the naïve CD4T-cell subfraction, as estimated using EpiDISH.

## Discussion

*OmniAge*, written in both R and Python, offers a comprehensive package of AOBs comprising over twice as many AOBs compared to competing tools like BioLearn, methylCYPHER or pyaging. As shown here, this increase in AOBs is critical for novel discovery and hypotheses generation, underscoring *OmniAge’s* added value.

One important finding gained with *OmniAge* was the striking correlation of mitotic age clocks with DNAm proxies of clonal hematopoiesis. Although this correlation was strongest for ASXL1 and TET2 CH, it was also seen for DNMT3A. This correlation between mitotic age and CH makes biological sense, since subclonal expansions associated with specific somatic mutations are driven by increased proliferation and hence an increased mitotic age. Another key but unexpected finding was the relatively strong correlation of mitotic clocks with AdaptAge, but not DamAge. The consistency of this finding across multiple independent large cohorts as well as the robust observation that the correlation between mitotic age and AdaptAge age-acceleration is strongest for the older age-group, clearly points towards a real biological effect. Of note, the AdaptAge causal clock was originally proposed to capture beneficial adaptations and hence the expectation would be that this clock should correlate with favorable outcomes ^11^. However, a recent assessment of 14 epigenetic clocks and 174 incident disease outcomes in the large Generation Scotland cohort showed that on average AdaptAge did not correlate with health outcomes, although it displayed large variance, being favourable for some outcomes and not for others ^71^. This ambiguity was also seen in the EPIC-Italy cohort, with AdaptAge age-acceleration predicting an increased risk of breast cancer but a lower risk of colon cancer. The unexpected positive correlation of AdaptAge with breast cancer risk not only highlights the underlying complexity of the AOB landscape, but also that an increased mitotic age in blood, as predicted with stemTOC, can be associated with the future risk of a solid cancer-type. It is also noteworthy that whilst a mitotic clock like stemTOC displayed strong correlations with AdaptAge and CH-proxies, neither AdaptAge nor DamAge associated with CH-proxies. Interestingly, although CH has generally been associated with adverse health outcomes ^48–50,72,73^, there is emerging evidence that many other forms of CH exist and that some of these could reflect beneficial adaptations ^51,74^. Thus, AdaptAge and DamAge may correlate with other forms of CH not considered here. In the context of the EPIC-Italy cohort, there were also key differences between cancer-types. Whilst TET2 and ASXL1-CH were associated with higher risk of developing breast cancer, they did not associate with risk of developing colon cancer. In contrast, DNMT3A-CH was strongly associated with risk of colon cancer but not breast cancer. Resolving why these patterns for causal clocks, CH-proxies and mitotic clocks are so different between breast and colon cancer risk is an important question for future research.

More generally, the patterns of correlation displayed by these AOBs with Episcores ^33^ (DNAm-based proxies of serum protein levels) also led to some important novel observations. Notably, whilst Episcores displayed similar correlative patterns with mitotic clocks and CH-proxies, specially ASXL1-CH ^75^, the Episcores associating most strongly with DamAge and AdaptAge were clearly distinct. In particular, unsupervised clustering revealed 3 broad clusters, with two antagonistic clusters displaying strong positive and negative correlations with mitotic clocks and CH-proxies, respectively. The contrast between these two clusters was striking: the cluster correlating positively involved chronic inflammation markers (macrophage chemokines like CCL17, CCL18, CCL22 and CRP), whilst the negatively correlated one appears to represent acute inflammation (MPO, S100A9). The cluster correlating positively with mitotic clocks and CH-proxies was further characterized by ECM degradation & tissue remodelling (MMPs, FAP) as well as chronic endothelial dysfunction (ESM-1, Semaphorin-3E), whilst the negatively correlated cluster contained markers of acute endothelial activation (e.g. E-selectin). Further underscoring this dichotomy, positively correlated Episcores comprised markers of myeloid skewing and immuno-senescence (G-CSF, CCL11, CD163), whilst anti-correlated Episcores like TNFRSF17 mark lymphoid adaptive function. Of note, the association between CH mutations like TET2, DNMT3A and ASXL1 with myeloid skewing is well-known ^76,77^. Thus, whilst the positively correlated Episcores appear to capture chronic systemic effects and myeloid lineage skewing, the anti-correlated ones appear to capture more acute and transient processes that are often related to specific tissue functions (liver: HGFA, neuron: Contactin-4, bone: osteomodulin). Hence, this suggests that epigenetic mitotic clocks and the ASXL1/TET2/DNMT3A CH-proxies are tracking a shift from a responsive, adaptive, and acutely reactive physiology to a state of chronic, myeloid-driven, tissue-degrading systemic dysregulation.

Episcores also displayed a striking bi-modal association with causal clocks, with those correlating most positively with DamAge simultaneously displaying strong anti-correlations with AdaptAge. Focusing on the 3 Episcores (NOTCH1, GDF-8 and CD209) displaying strongest anti-correlations with DamAge, it is striking that these proteins are implicated in tissue regeneration (NOTCH1) ^55–57^, regulation of skeletal muscle mass (GDF-8) ^58–60^, and dendritic immune surveillance (CD209) ^78,79^, respectively. These represent important biological processes that, if disrupted, can accelerate aging. This makes them not only interesting aging biomarkers but also highlights the NOTCH pathway, muscle-immune crosstalk, and dendritic cell biology as critical areas for putative interventions aimed at reducing causal epigenetic damage.

It is also noteworthy that the correlation between mitotic age and AdaptAge age acceleration was much stronger for the mitotic clocks based on gains of DNAm (e.g. epiTOC1, stemTOC), compared to those based on loss of DNAm (e.g. RepliTali). A potential explanation for this could be that the CpGs entering mitotic clocks like RepliTali or Hypoclock are more likely to be immune cell-type specific, which can confound mitotic age estimates with underlying changes in immune cell-type composition, as pointed out by us previously ^20,22^. Thus, the stronger correlation of epiTOC1 and stemTOC with AdaptAge age acceleration may well indicate the increased accuracy of these clocks to measure mitotic age, consistent with our previous findings ^20,22^.

Another interesting observation was the strong association of stochastic clocks with CH proxies. Overall, these stochastic clocks displayed significantly higher correlations with CH than the original clocks (Zhang, Horvath, PhenoAge) they derive from. This is surprising because these clocks and their stochastic counterparts are defined over the exact same CpGs. Hence, our finding suggests that DNAm-changes associated with CH, once constrained to susceptible genomic regions, are to a large extent stochastic and that 1^st^ generation clocks like Zhang and Horvath are capturing, to some degree, effects of CH. This is entirely plausible since CH is an ubiquitous highly reproducible feature of an aging immune system that would have been captured by any machine-learning algorithm aiming to predict chronological age. Moreover, in the case of Horvath’s pan-tissue clock, which was also trained on solid tissues, it is worth recalling that many solid tissues do display significant immune-cell infiltration ^80,81^. Hence, CH-signatures in blood are likely to be also present in the solid tissues used to train Horvath’s clock. It follows that many of the CpGs entering the Zhang and Horvath clocks are likely to be measuring this age-related increase in CH. Moreover, that the associated DNAm changes would be largely stochastic is consistent with the loss of epigenetic fidelity and maintenance following a mutation in an epigenetic enzyme like TET2 or DNMT3A. This contrasts with more deterministic DNAm changes, as observed for instance in relation to specific exposures like smoking, where the exact same CpGs mapping to say the AHRR locus are altered in the majority of exposed individuals ^47,82,83^.

Another appealing feature of *OmniAge* is the inclusion of molecular clocks representing other data modalities like transcriptomics. This inclusion facilitates direct comparison of different clocks on multi-modal datasets, such as the mRNA and DNAm dataset of the MESA study ^66^. As shown here, by applying a pan-tissue transcriptomic clock and various DNAm-clocks to the MESA dataset, age-acceleration estimates derived from different data modalities do in fact correlate, although this correlation was only prominent for CD4+ T-cells, with the association in monocytes being marginal and restricted to the PhenoAge clock. That the strong correlation in CD4+ T-cells can be attributed to underlying shifts between the naïve and memory CD4+ T-cell subsets highlights once again the importance of cell-type heterogeneity when deriving and interpreting transcriptomic or epigenetic clock estimates ^4^. Indeed, upon adjustment for the naïve CD4+ T-cell fraction, the correlation between transcriptomic and DNAm-clocks was strongly attenuated, suggesting that once we adjust for cell-type heterogeneity, clocks from different data modalities are not correlated or only marginally so, consistent with recent reports ^84,85^.

Finally, we acknowledge some limitations. Whilst *OmniAge* is a hypothesis generation and discovery tool, it only computes the DNAm-ages or scores for a large number of AOBs, leaving the subsequent hypothesis generation or discovery task to the user. We envisage that OmniAge’s usefulness will increase if it could be integrated with agentic expert systems, including for instance ClockBase ^86^. Second, whilst we have showcased many examples where *OmniAge* has led to novel findings, these merit further in-depth investigation to better explain their mechanistic underpinnings.

In summary, we envisage that *OmniAge’s* flexibility to estimate biological age across hundreds of AOBs and in two different programming languages (R & Python) will make this a popular application and hypothesis-generating tool in the aging field. *OmniAge* R and Python versions are freely available from https://github.com/Duzhaozhen/OmniAge. For users without programming experience, a webserver implementing OmniAge is available from https://www.biosino.org/omniage.

## Methods

### Implementation and quality control

We curated a library of 413 aging ‘omic’ biomarkers (AOBs) within the OmniAge framework, encompassing 12 categories that range from traditional epigenetic clocks to cross-species support (see **SI table S1** for a complete detailed listing). All algorithms were implemented in both R and Python environments. A strict quality control protocol was applied to validate the implementation of each AOB. We systematically compared the predictions generated by OmniAge with those obtained from the original software released with the respective publications. For biomarkers defined solely by published coefficients (i.e., where original code was unavailable), we validated our implementation against established third-party tools, including *pyaging* ^39^, *methylCIPHER* ^40^, and *methylclock* ^87^, to ensure high concordance and reliability.

### Blood-based DNA Methylation Datasets

In this study, we analyzed DNA methylation (DNAm) profiles from multiple independent blood-derived cohorts:

#### NSPT & TZH cohorts

The NSPT dataset ^62^ comprises whole blood samples from Han Chinese individuals, assayed using the Illumina Infinium MethylationEPICv1 platform. Data preprocessing and normalization were performed as previously described by Peng et al ^62^. Following quality control, the final dataset consisted of 811,870 probes across 3,501 samples. The study population had a mean age of 50 years (*-*SD ± 13; range: 17–83 years). A subset of the NSPT cohort, consisting of 705 samples, all from the same batch, is referred to as the ‘TZH’ cohort ^88^.

#### HPT-EPIC cohort

This dataset consists of peripheral blood samples from African-American participants in the Genetic Epidemiology Network of Arteriopathy (GENOA) study ^89^(GEO accession: GSE210255). Raw IDAT files were preprocessed using the *minfi* R package ^90^. Probes with a detection P-value >= 0.05 were filtered out. The data were subsequently normalized using the BMIQ method ^91^ to correct for type-2 probe bias. This yielded a normalized DNAm matrix encompassing 826,512 CpGs and 1,394 samples.

#### Johansson, Hannum, and Airwave cohorts

The Johansson ^42^ (729 whole blood samples, White Caucasians, Illumina 450k), Hannum ^92^ (656 whole blood samples, Whites, African Americans and Hispanics, Illumina 450k) and Airwave ^61^ (1032 PBMC samples, White Caucasians, Illumina EPICv1) datasets were downloaded and normalized as described previously by us ^61^.

#### EPIC-Italy

The European Prospective Investigation into Cancer and Nutrition (EPIC) is an ongoing study designed to investigate diet, nutrition, lifestyle and environmental factors with respect to cancer incidence. The cohort consists of 519,978 participants from 23 centres in 10 European countries - Denmark, France, Germany, Greece, Italy, the Netherlands, Norway, Spain, Sweden and the United Kingdom. Information on lifestyle, diet, anthropometric measures and environmental exposures were collected using questionnaires at recruitment and were standardized across the different participating centres ^93^. Blood was also collected from the majority of subjects at recruitment. Here we focused on the Italy subcohort, for which Illumina 450K data is freely available from the NCBI GEO website under the accession number GSE51032 (https://www.ncbi.nlm.nih.gov/geo/query/acc.cgi?acc=GSE51032). The file “GSE51032_RAW.tar” which contains the IDAT files was downloaded and processed with minfi package ^90^. Probes with NAs (defined by P > 0.01) were discarded. The filtered data was subsequently normalized with BMIQ ^91^, resulting in a normalized data matrix for 432,478 probes and 845 samples. Of these 845 samples, at last follow-up (2010), 424 remained cancer-free, with the rest having developed breast cancer (n=235: 233 women, 2 men), colon cancer (n=139), rectal cancer (n=12), skin cancer (n=4), rectosigmoid junction cancer (n=15), bladder cancer (n=5), prostate cancer (n=4), lung cancer (n=2) and other various forms of cancer (n=5).

#### Rheumatoid Arthritis EWAS (LiuRA)

This Illumina 450k whole blood dataset derives from Liu et al ^65^ and is available from GEO (GSE42861). It consists of 354 Rheumatoid Arthritis cases and 335 controls, with an age range from 18 to 70. We used the processed and normalized data as described in our previous publication ^44^.

#### Lehne cohort

This 450k DNAm dataset comprises over 2,700 peripheral blood DNAm profiles ^94^. In this study, we utilized the QC-processed and normalized version of this dataset as described by Voisin et al ^95^.

### MESA: mRNA expression and DNAm of sorted CD4+ T-cells and CD14+ monocytes

The Multi-Ethnic Study of Atherosclerosis (MESA) study performed mRNA expression (Illumina HumanHT-12 V4.0 expression beadchip) and DNAm (Illumina 450k) profiling in 1202 classical monocytes and 214 CD4+ T-cells. Data is publicly available from GEO under accession number GSE56047. DNAm data was processed as described by us previously ^20^. In particular, minfi and BMIQ-normalized DNAm data was adjusted for a covariate representing race, sex and batch. Raw data were normalized via the neqc algorithm within the limma package ^96^, which performed background correction, quantile normalization, and log2 transformation. This preprocessing resulted in yielding 17,377 genes (Ensembl Gene IDs) for downstream analysis.

### Single-cell RNA-Seq and sorted immune cell mRNA expression datasets

We downloaded the Seurat objects of two of the largest scRNA-Seq datasets of peripheral blood mononuclear cells (PBMCs) ^67,68^. For both datasets, we used the log-normalized data: for Yazar et al this was defined over 36571 genes (Ensembl Gene IDs) and 1,248,980 PBMCs from 981 donors with age-range (19, 97). For AIDA, it was defined over 36406 genes (Ensembl Gene IDs), 1,265,624 PBMCs and 625 donors with age range (19, 77). For our purposes we focused on donors for which there were at least 100 naïve CD4+ T-cells, and at least 100 memory CD4+ T-cells (combining effector and central memory ones). This left us with 844 and 497 donors for Yazar and AIDA, respectively. PASTA was then applied to pseudo bulk averages for each of the naïve and memory CD4+ T-cell subsets in each donor. PASTA age-acceleration was defined by regressing the clock estimates against chronological age, and these age-acceleration estimates were compared between CD4+ T-cell subtypes across donors using a one-tailed paired Wilcoxon rank sum test.

We also downloaded two microarray gene expression datasets profiling sorted CD4+ T-cell naïve and memory subtypes from GEO (GSE76598 and GSE94510). The GSE76598 dataset consists of 6 naive CD4T and 6 memory CD4T cell samples, sorted according to CD4+CD25-CD45RA+ and CD4+CD25-CD45RA-, respectively ^97^. Gene expression data was generated using Affymetrix Human Genome U133A 2.0 Array. Data was normalized using Robust Multiarray Average (RMA) and quantile normalization ^96^, leaving a total of 12650 genes. The second dataset (GSE94510) profiled 6 naive CD4T, 6 central memory CD4T and 6 effector memory CD4T cell samples, sorted according to CD4+CD45RO-CCR7+CD95-, CD4+CD45RO+CCR7+ and CD4+CD45RO+CCR7-, respectively ^98^. Data was generated using Affymetrix Human Genome U133 Plus 2.0 Array, and was normalized according using frozen RMA (fRMA) and quantile normalization, leaving a total of 19439 genes. Although age-information for both microarray sets was not available, all cell-types are derived from the same 6 donors and hence are age-matched, allowing comparison of age-acceleration estimates. Differences in PASTA estimates between cell-types was done using either a one-tailed paired Wilcoxon test (GSE76598) or a linear regression with donorID as a covariate (GSE94510).

### DNAm-based CH-proxies

We computed DNA methylation proxies for Clonal Hematopoiesis of Indeterminate Potential (CHIP), targeting four distinct categories: composite CHIP (’Any CHIP’) and gene-specific mutations in *DNMT3A*, *TET2*, and *ASXL1*. The specific CpG signatures and their corresponding directional effects were defined based on the meta-analysis coefficients derived from a recent large-scale multi-ethnic Epigenome-Wide Association Study (EWAS) ^35^. Specifically, the sign of the fixed-effect coefficient determined whether a CpG was classified as positively or negatively associated with CHIP status. To calculate the proxy for each CHIP category, we first identified the intersection of CpGs between the reference signature and the target dataset. For each common probe, methylation values were standardized to Z-scores across all samples to ensure a mean of zero and a standard deviation of one. The final CH-proxy for each individual was defined as the average Z-score of positively associated CpGs minus the average Z-score of negatively associated CpGs.

### DNAm-based CRP-scores

We implemented two DNA methylation-based scores to estimate serum C-reactive protein (CRP) levels. First, we computed the hsCRP score based on a signature of 1,765 CpGs identified in a large multi-ethnic meta-analysis (n>20,000) ^54^. Notably, while the identification of these CpGs included adjustment for six major immune cell fractions (granulocytes, monocytes, CD4+ and CD8+ T-cells, B-cells, and NK-cells), the analysis did not account for variation between naïve and memory lymphocyte subsets. The score is defined as the Pearson correlation coefficient between a sample’s Z-score normalized methylation profile and the directional signature (+1/−1) of these CpGs.

Second, to mitigate potential confounding by lymphocyte differentiation states (specifically naïve versus memory subsets), we derived a refined metric termed the intrinsic hsCRP score. We re-analyzed sorted immune-cell DNAm data ^61,70^ to identify CpGs within the original signature that are stable across differentiation. We selected 62 CpGs that showed no significant methylation differences (nominal limma P > 0.2) between naïve and memory subtypes across B cells, CD4+ T cells, and CD8+ T cells. The intrinsic hsCRP (intCRP) score is calculated using the same Pearson correlation framework restricted to this 62-CpG subset.

### Correlation and clustering analysis of AOBs

Prior to correlation analysis, we performed strict quality control to ensure biomarker compatibility. We restricted the analysis exclusively to AOBs compatible with Illumina 450K/EPIC array platforms and derived from whole blood tissue. Subsequently, we calculated age-acceleration residuals for each biomarker to remove confounding effects: for the Johansson and TZH cohorts, we regressed each AOB against chronological age and sex (AOB∼Age + Sex). For the Hannum cohort, to account for its multi-ethnic composition (Whites and Hispanics), we additionally included ethnicity as a covariate in the model (AOB∼Age + Sex + Ethnicity). To explore the global relationship between biomarkers we calculated pairwise Pearson correlation coefficients between the residuals of all AOBs. To investigate the intrinsic latent structure beyond simple pairwise associations, we performed unsupervised clustering using a model-based approach. Specifically, the residuals were Z-score standardized and clustered using a Gaussian finite mixture model implemented in the *mclust* R package(v6.1.1) ^99^. In the Johansson dataset, we selected the optimal model parametrization and number of clusters (k) based on the Bayesian Information Criterion (BIC). To facilitate comparability with the clusterings in Hannum, TZH and Airwave cohorts, we ran *mclust* on these cohorts using the optimal *k* as inferred in Johansson. This was done, because in the TZH cohort, the BIC failed to predict the correct number of clusters due to the EM-algorithm converging on a low-cluster number solution with clusters defined by abnormally high variance (a well-known problem with EM-optimization).

### Construction and validation of the immune cell-type fraction clock

We utilized the NSPT cohort ^62^ (n=3,501) to train the clocks based on 12 immune cell-type fractions estimated via EpiDISH ^61,63^. Two algorithms were employed: (1) Random Forest: Implemented using the *randomForest* R package (v4.7-1.2) ^100^. We performed a systematic evaluation to optimize the number of trees (ntree) using ten-fold cross-validation evaluating values ranging from 50 to 800 in increments of 50. The optimal ntree was identified as the value that minimized the cross-validated Mean Absolute Error (MAE). The final model was constructed using this optimized parameter with a minimum node size of five. (2) Lasso Regression: Implemented using the *glmnet* R package (v4.1.8) ^101^ with the elastic net mixing parameter a = 1. The optimal penalty parameter A was selected via ten-fold cross-validation to minimize prediction error. Both models were subsequently validated in four independent datasets: Johansson(450k, n=729) ^42^, Hannum(n=656) ^43^, Airwave(EPIC, n=1,032) ^61^, and HPT-EPIC(EPIC, n=1,394) ^89^.

### Cancer risk incidence analysis in EPIC-Italy

OmniAge was applied to the BMIQ normalized DNAm dataset of EPIC-Italy cohort. Correlations between AOBs were computed across all 845 blood samples. For cancer risk incidence analysis we restricted to two subcohorts. One consisting of 233 women who developed breast cancer after sample draw (cases) and 340 women who did not develop any cancer (controls). Among the 233 breast cancer cases, the average time to diagnosis was 5.36 +/− 3.69 years, IQR=1.91-8.59, with maximum of 14.1 years. The other subcohort consisted of 139 individuals (both sexes) who developed colon cancer after sample draw (cases) and 424 individuals who did not develop any cancer (controls). Among the 139 colon cancer cases, the average time between sample collection to diagnosis was 6.38 +/− 3.55 years, IQR=3.38-8.86, with maximum at 14.4 years. AOBs were standardized to unit variance and correlated to breast or colon cancer incidence using Cox proportional hazards regressions. We performed both univariate regressions as well as multivariate regressions adjusting for chronological age and sex (in the case of colon cancer). In addition, we estimated the fractions of 12 immune-cell types using EpiDISH ^61,63^, which were included as covariates alongside age and sex in a second multivariate analyses, to determine if any residual associations with cancer risk are driven by variations in immune-cell fractions.

### Cross-correlation of Episcores with mitotic clocks, causal clocks and CH-proxies

Using the EPIC-Italy cohort (n=845 blood samples) we computed Pearson correlations (PCCs) of 111 Episcores (109 Episcores from Gadd et al ^33^ plus a CRP-proxy ^54^ (described earlier) and another one for IL-6 levels ^32^) with 4 mitotic clocks (stemTOC ^22^, epiTOC1^18^, RepliTali ^21^ and HypoClock ^19^), causal clocks AdaptAge and DamAge ^11^ and the 4 CH-proxies described earlier. Pearson correlations were computed after adjusting these 10 AOBs for age and sex. Unsupervised clustering of PCCs was performed using an Euclidean distance metric. When ranking Episcores with the 10 AOBs, in the case of mitotic clocks, we removed HypoClock, because this is known to be confounded by variations in immune-cell fractions ^20^. Of note, the original 109 Episcores were derived from two different proteomic platforms (SomaScan + Olink) but a few proteins measured in both (e.g. GZMA).

### Application of transcriptomic clocks and DNAm clocks to joint mRNA-DNAm MESA dataset

We applied various transcriptomic and DNAm clocks to the normalized mRNA expression and DNAm datasets of sorted monocytes (n=1202) and CD4+ T-cells (n=214) from the MESA study ^66^. Specifically, the cell-type-specific transcriptomic clocks, collectively termed scImmuAging ^27^, were applied in a matched manner: the scImmuAging CD4+ T-cell clock was evaluated exclusively on the CD4+ T-cell transcriptomic data, while the scImmuAging monocyte clock was applied to the monocyte data. In contrast, the pan-tissue transcriptomic clock PASTA ^26^ and all evaluated DNAm clocks were applied to the datasets of both cell types. For all clocks, Residual Age Acceleration (RAA) was calculated as the residuals from a linear regression model fitting the clock-estimated biological age against chronological age and a composite categorical covariate encompassing all unique combinations of race, sex, and batch. To explore whether the observed correlations between the pan-tissue transcriptomic clock (PASTA) RAA and the DNAm clock RAAs were driven by underlying cell-type heterogeneity, we performed an additional analysis within the CD4+ T-cell dataset. Specifically, we used EpiDISH to estimate the relative proportions of naïve versus memory CD4+ T-cell subfractions. The estimated naïve CD4+ T-cell fraction was then included as an additional covariate when recomputing the RAA of the DNAm clocks.

### Application of PASTA to scRNA-Seq datasets and bulk mRNA expression of sorted cells

To further demonstrate that PASTA clock estimates differ between naïve and memory CD4+ T-cells of the same chronological age, we applied PASTA to two of the largest single-cell RNA-Seq datasets of peripheral blood mononuclear cells (Yazar et al ^67^ and AIDA ^68^). PASTA was applied to pseudo bulk averages for each of the naïve and memory CD4+ T-cell subsets in each donor (thus age-matched). PASTA age-acceleration was defined by regressing the clock estimates against chronological age, and these age-acceleration estimates were compared between CD4+ T-cell subtypes across donors using a one-tailed paired Wilcoxon rank sum test. In the case of the two microarray mRNA expression datasets of sorted CD4+ T-cell subtypes, age information of donors was not available, but all cell-type samples derived from the same 6 donors and hence are age-matched, allowing differential comparison of age-acceleration (clock) estimates. Differences in PASTA estimates between cell-types was done using either a one-tailed paired Wilcoxon test (GSE76598) or a linear regression with donorID as a covariate (GSE94510).

## Data availability

The primary datasets used in this study are all publicly available. The DNAm datasets of HPT-EPIC ^89^, Johansson ^42^, Airwave ^64^, Hannum ^92^, EPIC-Italy ^52^, Rheumatoid Arthritis EWAS ^65^ and Lehne ^94^ cohorts can be accessed via the NCBI Gene Expression Omnibus (GEO) using accession numbers: GSE210255 (HPT-EPIC, https://www.ncbi.nlm.nih.gov/geo/query/acc.cgi?acc=GSE210255), GSE87571 (Johansson, https://www.ncbi.nlm.nih.gov/geo/query/acc.cgi?acc=GSE87571), GSE147740 (Airwave, https://www.ncbi.nlm.nih.gov/geo/query/acc.cgi?acc=GSE147740), GSE40279 (Hannum, https://www.ncbi.nlm.nih.gov/geo/query/acc.cgi?acc=GSE40279), GSE51032 (EPIC-Italy, https://www.ncbi.nlm.nih.gov/geo/query/acc.cgi?acc=GSE51032), GSE42861 (Rheumatoid Arthritis EWAS, https://www.ncbi.nlm.nih.gov/geo/query/acc.cgi?acc=%20GSE42861) and GSE55763 (Lehne, https://www.ncbi.nlm.nih.gov/geo/query/acc.cgi?acc=GSE55763). The Illumina EPIC DNAm data for NSPT cohort ^62^ can be requested at https://ngdc.cncb.ac.cn/omix/release/OMIX004363. The Illumina EPIC DNAm data for the TZH cohort ^88^. are deposited in the National Omics Data Encyclopedia (NODE) under accession OEP000260 and can be accessed by submitting a data access request (https://www.biosino.org/node/project/detail/OEP000260). The DNAm and mRNA expression data from the MESA study is available from GEO under accession number GSE56047 (https://www.ncbi.nlm.nih.gov/geo/query/acc.cgi?acc=GSE56047). The two microarray gene expression datasets profiling sorted CD4+ T-cell naïve and memory subtypes is available from GEO under accession number GSE76598 (https://www.ncbi.nlm.nih.gov/geo/query/acc.cgi?acc=GSE76598) and GSE94510 (https://www.ncbi.nlm.nih.gov/geo/query/acc.cgi?acc=GSE94510).

## Code availability

*OmniAge* is freely available as an R or Python package from https://github.com/Duzhaozhen/OmniAge. There we provide installation instructions and user-friendly tutorials. These tutorials for both R and Python versions are also available as additional Supplementary Information Files (OmniAgeR_tutorial.html & OmniAgePy_tutorial.html). The Python version of OmniAge is also available from https://pypi.org/project/omniage/.

## Acknowledgements

This work was supported by NSFC (National Science Foundation of China) grants, grant numbers W2431024, 32370699 and 32570775.

## Authors contributions

AET conceived the study. ZD performed the statistical and bioinformatic analyses and produced the R and Python versions of OmniAge. Manuscript was written by AET with contributions from ZD. YL helped with the webserver deployment. HT and XG contributed data and assisted with bioinformatic analyses.

